# Patterns of Mitochondrial ATP Predict Tissue Folding

**DOI:** 10.1101/2025.08.31.673364

**Authors:** Bezia Lemma, Megan Rothstein, Pengfei Zhang, Bridget Waas, Marcus Kilwein, Safiya Topiwala, Sherry X. Zhang, Anvitha Sudhakar, Katharine Goodwin, Elizabeth R. Gavis, Ricardo Mallarino, Andrej Kosmrlj, Celeste M. Nelson

## Abstract

The construction of complex tissue shapes during embryonic development results from spatial patterns of gene expression and mechanical forces fueled by chemical energy from ATP hydrolysis. We find that chemical energy is similarly patterned during morphogenesis. Specifically, mitochondria are locally enriched at the apical sides of epithelial cells during apical constriction, which is widely used across the animal kingdom to fold epithelial tissues. Timelapse imaging, spatial transcriptomics, and measurements of oxygen consumption rate reveal that mitochondrial density, potential, and ATP increase in epithelial cells prior to actomyosin contraction and tissue folding, which is prevented by inhibiting oxidative phosphorylation. Mitochondrial enrichment and apicobasal patterning are conserved during apical constriction in flies, chicks, and mice, and these subcellular patterns can be used to predict computationally patterns of tissue folding. These findings highlight a spatial dimension of bioenergetics in embryonic development.

## Main Text

Morphogenesis is a physical process in which tissues change shape in response to spatiotemporal patterns of gene expression and mechanical force (*1, 2*). While it is axiomatic that these processes require chemical energy in the form of ATP, surprisingly little is known about the spatiotemporal regulation of energy production, particularly by mitochondria (*3*). Recent attention has focused on the appearance of glycolytic gradients in chicken and mouse embryos, including during delamination of neural crest cells (*4*), migration of epiblast cells into the primitive streak (*5*), and posterior elongation of the body axis (*6-8*). These gradients have been interpreted in the context of metabolic signaling, as through the Wnt pathway (*9*), rather than in the context of energetics, although it has been postulated that glycolytic gradients may provide ATP for actin polymerization and thus promote cell motility (*6, 10*). The direct role of energy metabolism in powering the fundamental morphogenetic processes that fold tissue shapes, especially through mitochondrial production of ATP, remains largely unexplored.

One such fundamental morphogenetic process is apical constriction, a conserved motif in which epithelial tissues fold when cells shrink their apical surfaces by actomyosin contraction (*11-13*). The molecular machinery that generates the forces necessary for apical constriction is also conserved: ATP hydrolysis leads to phosphorylation of the myosin regulatory light chain and promotes assembly and activity of myosin motors, which contract the actin network that is anchored by adherens junctions to the apical cortex of the cell. During this morphogenetic process, myosin phosphorylation and the generation of force are spatially patterned both within the cell (at the apical side) and within the tissue (in the actively contracting cells). That myosin motors require ATP hydrolysis prompts the question of whether mitochondrial energy metabolism is similarly spatially patterned.

To investigate the possible spatial relationship between mitochondria and generation of mechanical forces during apical constriction, we began by using the embryonic chicken lung as a model system. The early embryonic chicken lung consists of tubes of simple columnar epithelium surrounded by mesenchyme (*14*). The first three branches form at predictable locations and times along the dorsal surface of the primary bronchus (**Fig. S1a-c**) (*15-17*). Epithelial cells at these locations undergo apical constriction (**Fig. 1a**), which folds the epithelium into the adjacent mesenchyme and thus initiates a branch (*16*). We took advantage of this stereotypy to systematically visualize mitochondrial membrane density within the epithelium during morphogenesis. Specifically, we isolated embryonic lungs at timepoints in between initiation of the first branch (t=116 hr; Hamburger-Hamilton stage HH24) and initiation of the third branch (t=128 hr; HH27). As a metric for mitochondrial membrane density, we conducted immunofluorescence analysis for Tom20, an outer mitochondrial membrane import receptor subunit (**Fig. 1b**). This immunostaining revealed that mitochondrial membrane density is enriched in the epithelium within nascent branches. Counterstaining each sample for E-cadherin allowed us to generate three-dimensional (3D) segmentations of the epithelial tree to quantify Tom20 enrichment (**Fig. 1b, c; Movie S1**). Quantitative image analysis confirmed that mitochondrial membrane density is significantly higher in the epithelium than in the mesenchyme, and significantly higher in the medial epithelium that forms branches than in the proximal epithelium that does not (**Fig. 1d**). Consistently, unbiased spatial transcriptomics analysis revealed that mitochondrial gene expression is enriched within the branching epithelium (**Fig. S2a-f**). The spatial patterning of elevated mitochondrial density within the branching dorsal epithelium persisted when we normalized to the adjacent non-branching ventral epithelium across all stages examined (**Fig. 1e-f; Fig. S1d-f**). Curiously, we noticed a strong bias in mitochondrial localization within the branching epithelium, with elevated density on the apical side of the cells (**Fig. 1g, insets; Fig. S1g-i**), mirroring the patterns of actomyosin contraction at the apical surfaces of these regions of the tissue.

**Fig. 1.**
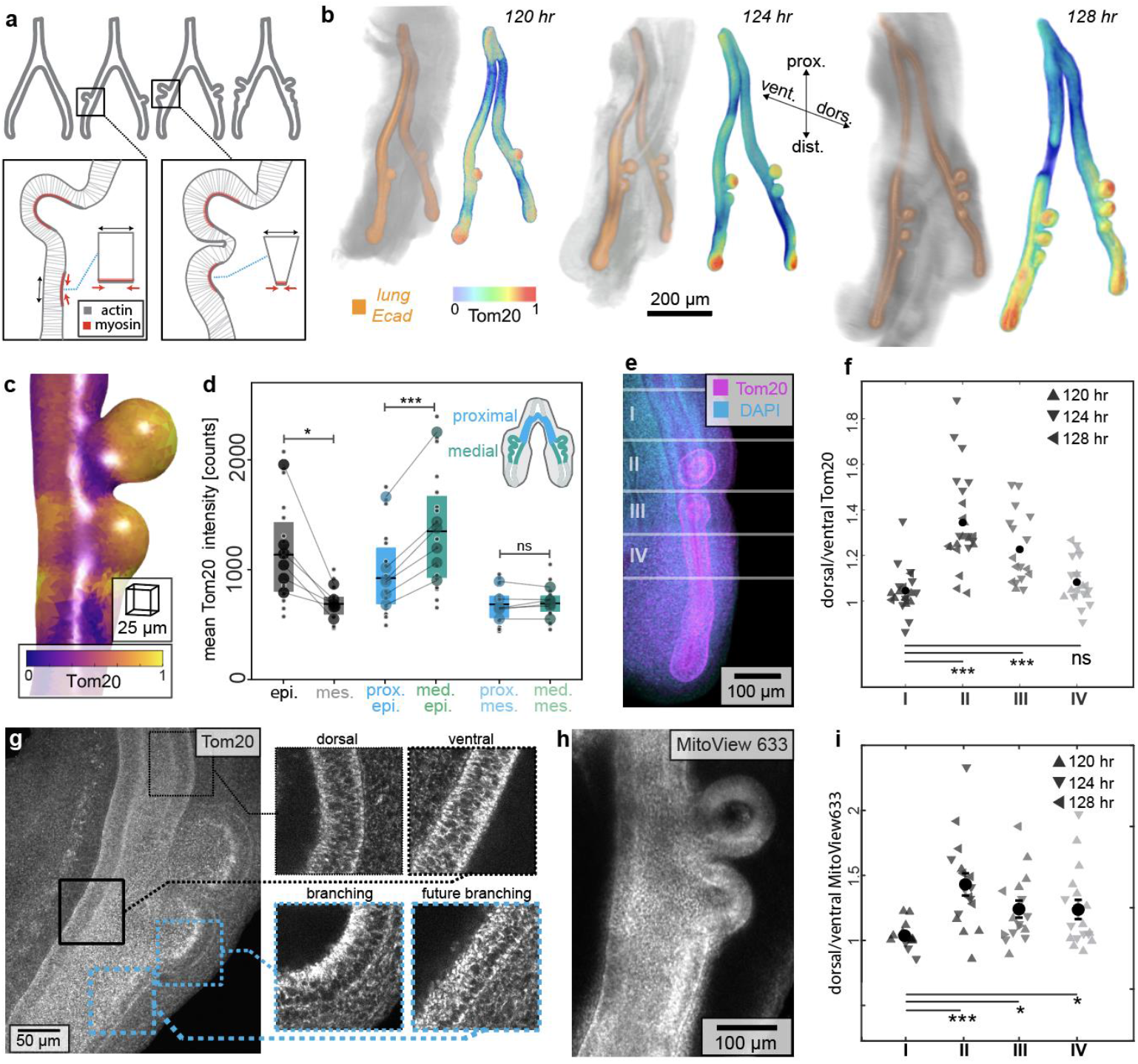
Mitochondrial membrane density and potential are locally enriched in the lung epithelium at branch sites prior to and during apical constriction. (**a**) Schematic of early embryonic chicken lung, indicating sites of apical constriction. (**b**) Projections of confocal images of staining for E-cadherin (orange, left) and Tom20 (color-coded, right) at different stages of development. Scale bar, 200 μm. (**c**) Local intensity of Tom20 within the epithelium, color-coded based on intensity. Scale bar, 25 μm. (**d**) Graph of Tom20 staining intensity in the entire epithelium and mesenchyme, proximal (non-branching) epithelium, medial (branching) epithelium, and adjacent mesenchymal regions. Shown are data from 18 lungs (small circles) across 6 independent replicates (means shown by large circles). Lines match independent replicates between measurements. Box plot shows median, first, and third quartile of data. *, *** indicate p=0.02, p=0.001 respectively. (**e**) Maximum-projection image of staining for Tom20 (magenta) and nuclei (blue), partitioned into four regions: (I) non-branching, (II) branching, (III) nascent branching, and (IV) future branching. Scale bar, 100 μm. (**f**) Graph of normalized Tom20 staining intensity in regions (I-IV) of lungs at different stages of development. Shown are data from 21 lungs across 9 independent replicates. *** indicates p=2e-7, p=9e-5 respectively (**g**) Maximum-intensity projection of staining for Tom20; insets show intensity and apicobasal patterning in single z-planes. Scale bar, 50 μm. (**h**) Maximum-intensity projection of MitoView633 fluorescence. Scale bar, 100 μm. (**i**) Graph of normalized MitoView633 intensity in regions (I-IV) of lungs at different stages of development. Shown are data from 17 lungs. ***, *, * indicate p=0.0003, p=0.01, and p=0.03 respectively.

To directly visualize mitochondrial energy metabolism, we took advantage of the fact that embryonic chicken lungs continue to undergo branching morphogenesis when cultured as explants (*15-18*). We isolated and cultured embryonic lungs in the presence of MitoView633, a mitochondrial membrane potential-dependent dye that accumulates at mitochondria and fluoresces in proportion to the voltage across the mitochondrial membrane (*19*) (**Fig. 1h**). As with staining for Tom20, we quantified the relative fluorescence intensity of MitoView633 as a function of position along the primary bronchus by normalizing to the adjacent ventral epithelium (**Fig. 1i**). We found that mitochondrial membrane potential is higher in the dorsal epithelium than in the non-branching ventral epithelium. Consistent with our analysis of mitochondrial membrane density and mitochondrial gene expression, we found that mitochondrial membrane potential is elevated at the apical sides of cells within branch sites in the epithelium, even prior to branch initiation (**Fig. 1h; Fig. S1j-m; Movie S2-S3**). These data are consistent with an increase in mitochondrial activity – and potentially higher levels of chemical energy in the form of ATP – within the epithelium as it undergoes apical constriction.

To determine directly whether sites of apical constriction correspond to elevated levels of ATP, we cultured lung explants in the presence of ATPRed1, which labels ATP surrounding mitochondria (*20*). We first calibrated the dye by measuring its fluorescence intensity as a function of ATP concentration (**Fig. S3a**). We also confirmed that blocking ATP synthesis by treating with the mitochondrial ATP synthase inhibitor, oligomycin, leads to a decrease in ATPRed1 fluorescence intensity (**Fig. S3b-d**). We then cultured embryonic lung explants in the presence of ATPRed1 during the branching process (**Fig. 2a; Movie S4**). We found that the fluorescence intensity of ATPRed1 is lowest in the non-branching epithelium, increases at future branch sites prior to apical constriction, and further increases within the epithelium of nascent branches (**Fig. 2b-c; Fig. S3e; Movie S5**). Blocking actomyosin contractility by treating explants with the myosin II inhibitor, blebbistatin, blocks both apical constriction as well as the formation of new branches without preventing epithelial growth (**Fig. 2d; Fig. S3f-g**) (*16, 21*). Curiously, we found that treatment with blebbistatin decreases mitochondrial ATP production, as inferred by quantifying the fluorescence intensity of ATPRed1 (**Fig. 2e-f; Movie S6**), but has no effect on mitochondrial membrane density, as inferred by quantifying Tom20 staining intensity (**Fig. 2g**). These data suggest that the process of apical constriction both activates and is coupled to mitochondrial ATP synthesis within the actively contracting region of the epithelium, even before detectable deformation of the tissue.

**Fig. 2.**
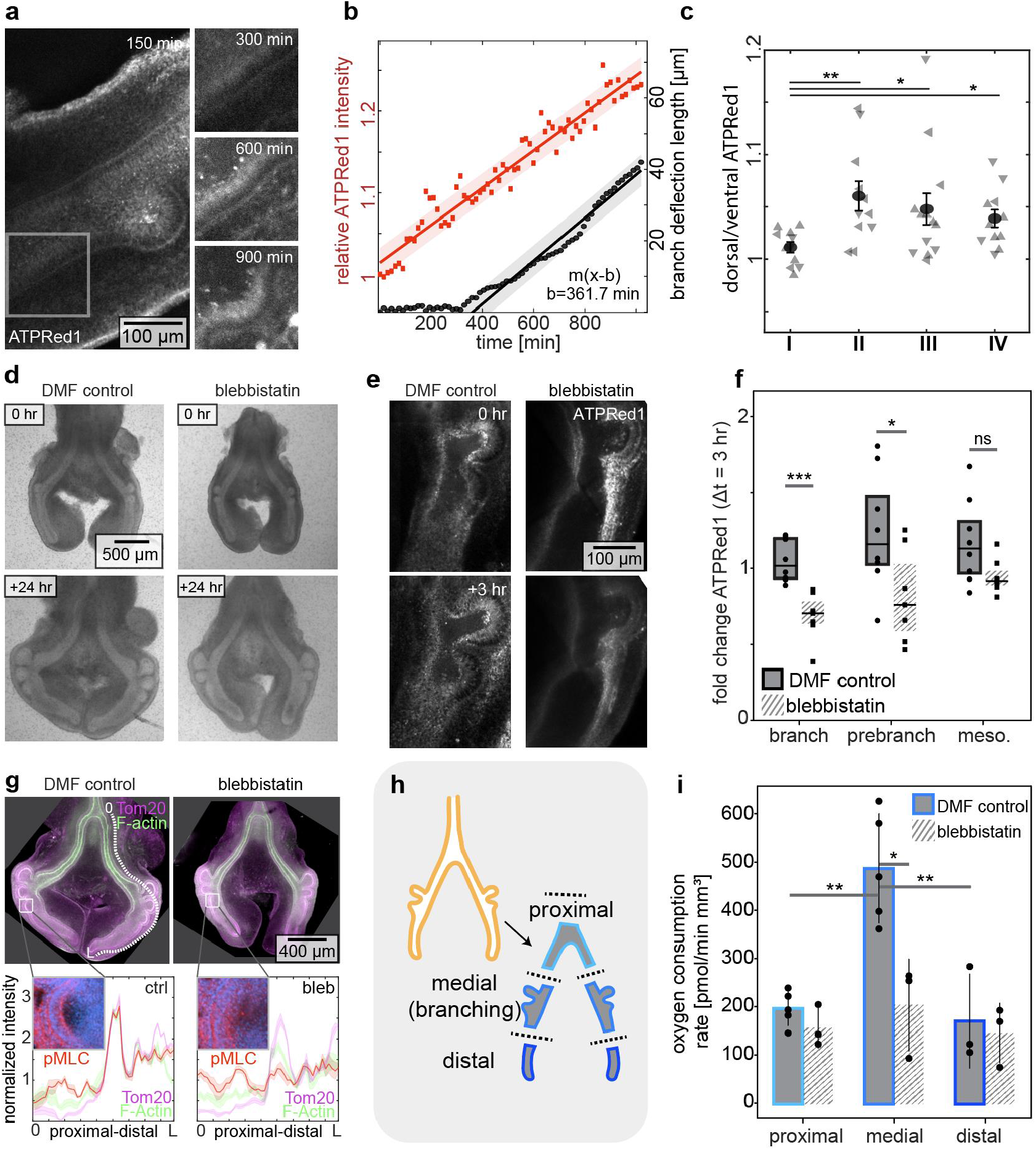
Mitochondrial ATP density and oxygen consumption rate are increased at sites of branch initiation both before apical constriction and in response to actomyosin contraction. (**a**) Timelapse images of ATPRed1 intensity during branch initiation in the embryonic chicken lung. Scale bar, 100 μm. (**b**) Graph showing ATPRed1 intensity at the dorsal side relative to the ventral side (red) and deflection of the epithelium (black) as a function of time, for the inset region shown in (a). Shaded band represents 95% confidence interval of linear fit. (**c**) Graph of normalized ATPRed1 intensity in regions (I-IV; defined in Fig. 1e) of lungs at different stages of development. Shown are data from 17 lungs. **, *, * indicate p=0.001, p=0.04, and p=0.02, respectively. (**d**) Phase-contrast images of lung explants before and 24 hours after culture in the presence of DMF control or blebbistatin (20 μM). Scale bar, 500 μm. (**e**) Maximum intensity projections of ATPRed1 staining immediately after and 3 hours after culture in the presence of DMF control or blebbistatin (20 μM). Scale bar, 100 μm. (**f**) Fold change in ATPRed1 over 2 hours of treatment in dorsal branching tissue. Shown are data from 8 independent control lungs and 7 independent blebbistatin-treated lungs. *, ** indicates p=0.03, and p=0.0004 respectively. (**g**) Top: maximum-intensity projection images of staining for Tom20 (magenta) and F-actin (green) in lungs cultured in the presence of DMF control or blebbistatin. Scale bar, 400 μm. Bottom: insets show pMLC (red) and nuclei (blue) at the region of an initiating branch. Graphs show pMLC from proximal (0) to distal (L) along the outer edge of the epithelium, as indicated by the dotted white line. Data are normalized and averaged over four samples. Shaded region represents standard error of the mean (s.e.m.). (**h**) Schematic of dissections used to divide lungs into proximal (non-branching), medial (branching), and distal (future branching) regions. (**i**) Graph showing oxygen consumption rate of proximal, medial, and distal regions of lungs cultured in the presence of DMF control or blebbistatin. Each dot indicates a replicate, with each replicate containing tissue pooled from 20 lungs. **, *, **, indicate p=0.003, p=0.013 and p=0.002, respectively.

Mitochondrial ATP synthesis relies on a proton gradient across the mitochondrial membrane, maintained by oxygen molecules accepting electrons at the end of the electron transport chain. Our data thus suggested that regions of the epithelium undergoing apical constriction might consume more oxygen than other regions of the tissue. To test this hypothesis, we surgically divided embryonic chicken lung explants into proximal (non-branching), medial (branching), and distal (future branching) regions (**Fig. 2h**), and then used Seahorse analysis to measure the oxygen consumption rate of each region separately (**Fig. S4a-i**). This analysis revealed a significantly higher oxygen consumption rate in the medial branching region of the lung, as compared to the proximal non-branching region (**Fig. 2i**), consistent with our hypothesis. As expected from our experiments using ATPRed1, we found that blocking apical constriction by treating explants with blebbistatin significantly attenuated the oxygen consumption rate of the branching medial region (**Fig. 2i**). Actomyosin-induced apical constriction thus coincides with and promotes mitochondrial respiration and synthesis of ATP.

Given that the patterns of mitochondrial density (Tom20 and spatial transcriptomics), membrane potential (MitoView633), ATP (ATPRed1), and oxygen consumption rate (Seahorse) either correspond to or presage those of apical constriction in the embryonic chicken lung epithelium, we hypothesized that mitochondrial respiration might be required for this developmental process (**Fig. 3a**). We tested this hypothesis by culturing embryonic lung explants in the presence of oligomycin, which halved the oxygen consumption rate within 2 hours after administration (**Fig. 3b; Fig. S4c**), consistent with a significant reduction in mitochondrial respiration. Strikingly, treatment with oligomycin completely blocked the initiation of new branches in the explants (**Fig. 3c, d; Movie S7**). We also found that treatment with oligomycin led to a reduction in the intensity of staining for both F-actin as well as phosphorylated myosin light chain (pMLC) at the apical surface of the epithelium (**Fig. 3e, f**). Importantly, blocking mitochondrial respiration had no effect on the spatial patterns of mitochondrial density: oligomycin-treated explants still showed elevated levels of Tom20 staining intensity at branch sites in the dorsal epithelium and on the apical sides of actively branching cells (**Fig. 3e-g**). Mitochondrial production of ATP is therefore required for the generation of new branches by apical constriction of the embryonic chicken lung epithelium.

**Fig. 3.**
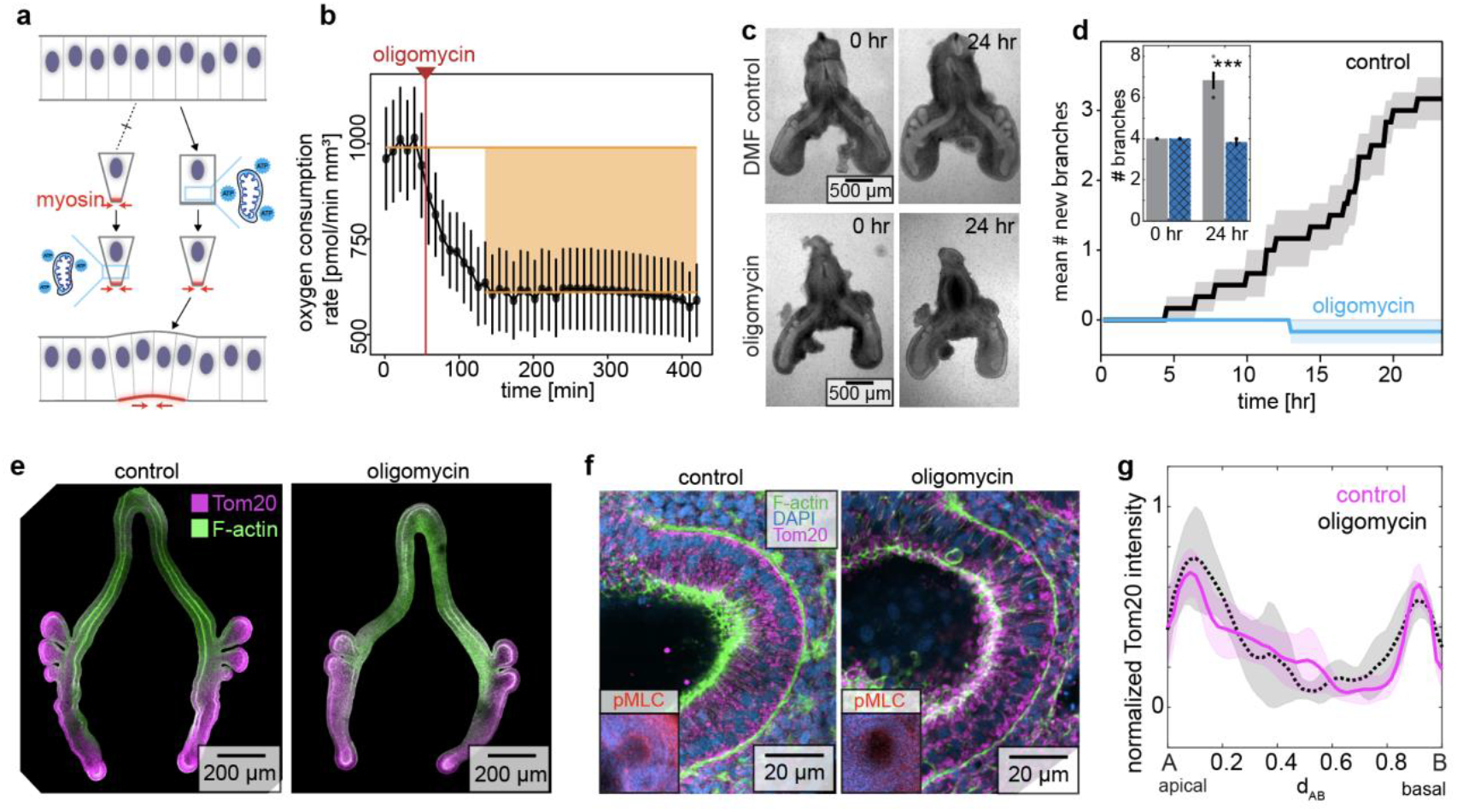
Mitochondrial ATP production is required for apical constriction and branch initiation in the embryonic chicken lung. (**a**) Schematic of two possible connections between mitochondrial energy and apical constriction. Actomyosin-driven apical constriction either precedes (left) or follows (right) mitochondrial ATP production. (**b**) Graph showing oxygen consumption rate of lungs as a function of time before and after treatment with oligomycin (2 μm). (**c**) Phase-contrast images of lung explants before and 24 hours after culture in the presence of DMF control or oligomycin (2 μm). Scale bars, 500 μm. (**d**) Graph showing the mean number of new branches as a function of time in lung explants cultured in the presence of DMF control or oligomycin (2 μm), averaged over 6 independent experiments. Shaded error bar indicates s.e.m. Inset bar graph shows initial and final number of branches. *** indicates p=0.000016. (**e**) Fluorescence images of staining for Tom20 (magenta) and F-actin (green) in lung explants cultured for 12 hours in the presence of DMF control or oligomycin (2 μm). Scale bars, 200 *μ*m. (**f**) Fluorescence images of staining for Tom20 (magenta), F-actin (green), and DAPI (blue). Insets show: DAPI (blue) and pMLC (red). Scale bars, 20 *μ*m. (**g**) Graph showing quantification of normalized Tom20 intensity as a function of position along the apicobasal axis of the branching epithelium, manually aligned for 3 independent staining experiments. Shaded error bar indicates s.e.m. at each position.

Apical constriction is a conserved morphogenetic motif that is used to fold simple epithelial sheets and tubes into more complex geometries, including during gastrulation (*22*), neurulation (*23*), and eye development (*24, 25*). To determine whether spatial patterns of mitochondrial density are conserved during apical constriction, we broadened our analysis across tissues and species. Specifically, we examined ventral furrow formation during gastrulation in *Drosophila* (**Fig. 4a**), neural tube closure during neurulation in chicken (**Fig. 4b**), and lens placode invagination during eye development in mouse embryos (**Fig. 4c**). Staining each system for Tom20 revealed spatial patterns of mitochondrial membrane density almost identical to those we observed in the chicken lung epithelium, with elevated intensity in the future ventral furrow (**Fig. 4d**), neural tube (**Fig. 4e**), and optic lens (**Fig. 4f**), and a bias to the apical side of the epithelium (**Fig. 4g-j**). Notably, mitochondrial accumulation in the ventral furrow aligns with previous observations of mitochondrial repositioning and fusion (*26*). Apical enrichment of mitochondria thus appears to be a defining feature of epithelial cells undergoing apical constriction.

**Fig. 4.**
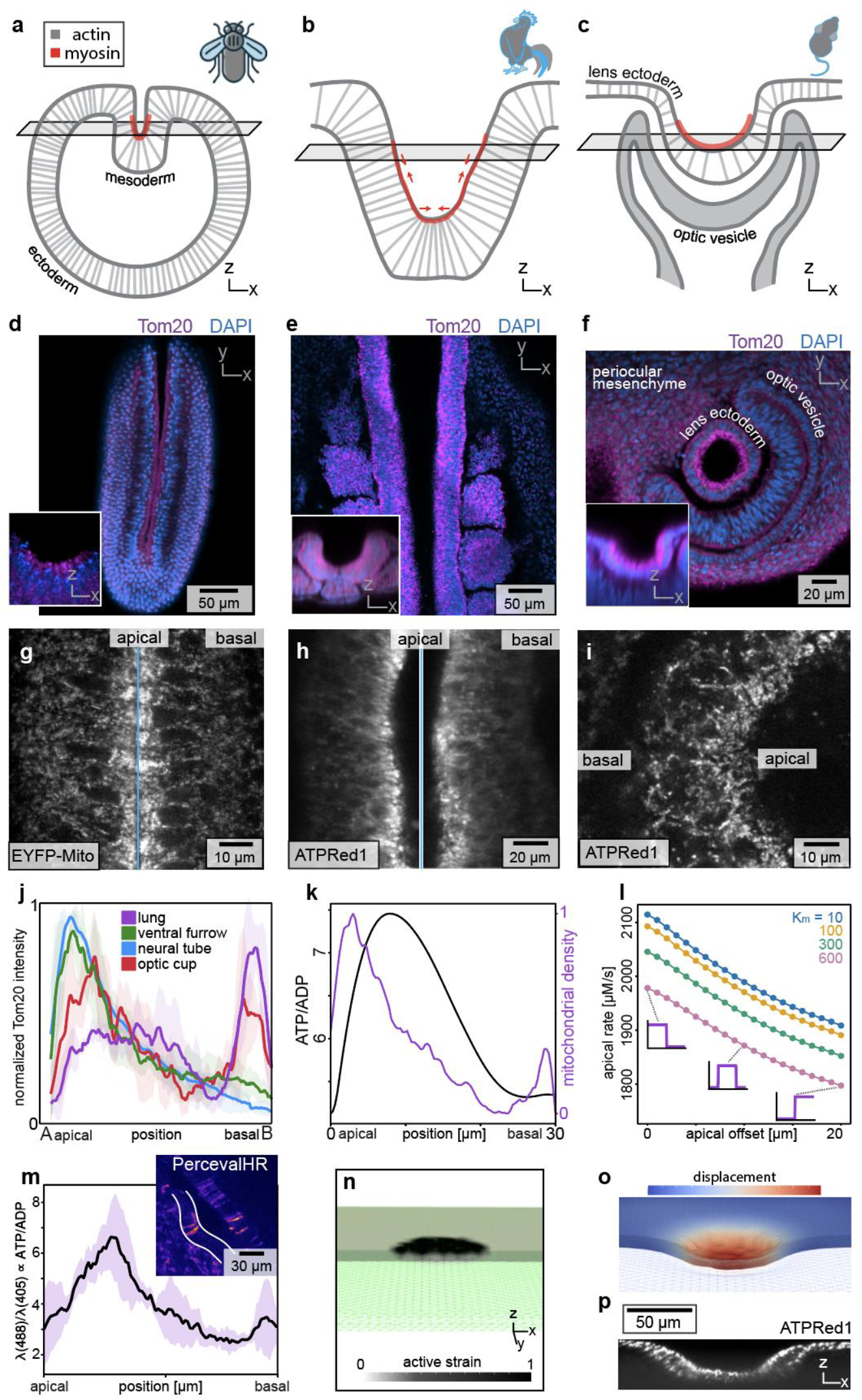
Patterns of mitochondrial membrane density are conserved and predict apical constriction across species and tissues. Schematics of (**a**) ventral furrow formation during gastrulation in *Drosophila*, (**b**) closure of the neural tube during neurulation in chicken, and (**c**) invagination of the lens placode during eye development in mouse embryos. (**d-f**) Confocal images showing staining for Tom20 (magenta) and nuclei (blue) during gastrulation, neurulation, and lens invagination. Insets show z-x sections. Scale bars 50 *μ*m, 50 *μ*m, and 20 *μ*m. (**g-i**) Confocal images showing live imaging of apicobasal mitochondrial patterning, as reported by EYFP-Mito in Drosophila, and ATPRed1 in chicken neural tube and mouse optic cup. Scale bars 10 *μ*m, 20 *μ*m, and 10 *μ*m. (**j**) Graph showing normalized Tom20 intensity in each tissue, revealing enrichment of mitochondria on the apical side. (**k**) Graph showing the ATP/ADP ratio (black) resulting from a mitochondrial density pattern (purple) in a 1D reaction-diffusion model, with Michaelis-Menten kinetics of hydrolysis on the apical side due to actomyosin in the region indicated in red and a baseline rate of hydrolysis throughout the cell. (**l**) Graph showing that the reaction-diffusion model predicts a decrease in myosin phosphorylation rate as the mitochondrial distribution is moved from the apical to the basal side of the cell, plotted for different values of the Michaelis constant *K_m_*. (**m**) Graph showing measurements of ATP/ADP ratio in cells undergoing apical constriction in branching epithelium of the chicken lung. Inset: confocal ratiometric image of PercevalHR. Scale bar 30 *μ*m. (**n**) Cutaway view of initial geometry for a simulation based on mitochondrial patterning of tissue deformation in the mouse optic cup; numerical mesh drawn in green, with applied active stress shown in grayscale. (**o**) Cutaway view of mesh deformed by active stresses, colored by the displacement field. (**p**) Confocal z-slice reconstruction of ATPRed1 signal in the mouse optic cup. Scale bar 50 *μ*m.

To determine whether the apicobasal patterning of mitochondrial density can be used to predict subcellular patterns of ATP concentration and, thus, actomyosin kinetics, we constructed a reaction-diffusion model of the cell. Specifically, we assumed that ADP is converted into ATP as a function of local mitochondrial density, at a rate determined by our Seahorse measurements, and hydrolyzed as a function of actomyosin and ATP concentrations, according to Michaelis-Menten kinetics. These mass-action kinetics predict a higher ATP/ADP ratio at the apical side of the cell along with a correspondingly higher rate of myosin phosphorylation (**Fig. 4k-l**). To test this prediction experimentally, we transduced embryonic chicken lungs with PercevalHR, a genetically encoded fluorescent biosensor that reports the intracellular ATP/ADP ratio (*27*). We then used this reporter to measure the ATP/ADP ratio during apical constriction and found, indeed, a significantly higher ratio at the apical side (**Fig. 4m**). To determine whether these subcellular patterns of mitochondria and rates of myosin phosphorylation might be sufficient to promote the observed multicellular patterns of folding, we then used the finite element method (FEM) to construct a 3D continuum model of each developing tissue, based on geometrical and biophysical parameters. Each tissue started as a simple shape (ellipsoid, half cylinder, or sheet, respectively), onto which we mapped patterns of apical constriction, as predicted by our observations of the subcellular patterns of mitochondria (**Fig. 4n; Fig. S5a-b**). Remarkably, each simulation produced a final geometry similar to that observed experimentally (**Fig. 4o; Fig. S5c-d; Movie S8**), suggesting that apicobasal patterns of mitochondrial density can be used to predict patterns of mechanical forces that drive changes in tissue shape. To determine whether we could similarly predict patterns of chemical energy within each developing system, we conducted timelapse imaging analysis of ATPRed1-labeled explants (**Fig. 4p**). In each case, we found significantly elevated fluorescence intensity in the regions of tissue predicted to undergo apical constriction (**Fig. S5e-j**). Collectively, our data suggest that the chemical energy from mitochondrial respiration is biased towards the apical sides of cells and coupled to the forces of tissue folding during apical constriction.

Mitochondrial density, membrane potential, and ATP production are elevated at the apical sides of cells before and during apical constriction. Our reaction-diffusion and FEM models suggest that the apical concentration of mitochondria would be sufficient to promote myosin phosphorylation at the apical cortex of the cell, thus promoting tissue folding. These observations, combined with previous studies focused on the molecular components of the force-generating machinery, suggest that the morphogenetic motif of apical constriction involves a conserved network in which spatially patterned energy metabolism is dynamically integrated with conserved genetic and mechanical components fold epithelia (*28, 29*). Importantly, we observe that the apicobasal patterning of mitochondrial density can be used to predict which cells within the epithelium will undergo apical constriction before appreciable changes in cell or tissue shape. Our findings suggest a new general paradigm: that subcellular patterns of organelles, energy, and energy metabolism are upstream of changes in tissue morphology during development. Consistent with this notion, waves of glucose uptake and glycolytic metabolism have been found to precede induction of epithelial-mesenchymal transition (EMT), a different morphogenetic motif that is important for development of the neural crest, primitive streak, and presomitic mesoderm (*4-10*), possibly downstream of signaling through fibroblast growth factor (FGF) (*6, 30, 31*). The upstream signals that promote the apical bias in mitochondrial localization prior to apical constriction remain to be uncovered.

Mitochondrial ATP production also appears to be essential for apical constriction, as treatment with oligomycin blocks actomyosin contraction. This observation could be taken to suggest that mitochondrial respiration is enhanced in these cells in lieu of glycolysis, as has been proposed for differentiating cells in the developing retina (*32*). However, our spatial transcriptomics and Seahorse data suggest that branching epithelia express high levels of glycolytic enzymes (**Fig. S2g**) and show elevated glycolytic extracellular acidification rates (**Fig. S4f, S4m-o**), suggesting a possible coupling between glucose metabolism and mitochondrial respiration, at least in the developing lung. While mitochondrial ATP is clearly necessary, quantifying the relationship between energy supply and mechanical work raises questions about efficiency. Estimations of the viscoelastic energy requirements necessary for apical constriction to deform the airway epithelium into a branch (**Methods** and **Movie S8**) suggest that the energy provided by mitochondrial respiration and glycolysis is staggeringly large (∼100,000,000 nJ) compared to the morphological deformation itself (∼10 nJ; **Methods**). A more accurate estimate will require teasing apart the energetic budget of the tissue and accounting for its energy demands beyond actomyosin contraction at the apical cortex. Nonetheless, this difference is vast, and might be explained by the low efficiencies with which energy is transduced into systematic molecular motion and force (*33, 34*). Alternatively, glycolysis and/or oxidative metabolism might promote biochemical signaling downstream of morphogens or reactive oxygen species to initiate apical constriction (*4, 35*). It will be intriguing to uncover both the upstream and downstream signals that link chemical energy, mechanical forces, and tissue folding.

## Abbreviations

3D: three-dimensional
FEM: finite element method
HH: Hamburger-Hamilton
pMLC: phosphorylated myosin light chain

## Acknowledgements

We thank the Tissue Morphodynamics Group for helpful discussions, G. Laevsky for assistance with imaging, D. Roman for electroporating glycolysis-reporter plasmids into embryonic lungs, and K. Zhang, I. Laderman, and R. Cui for testing new culture techniques.

## Author contributions

Conceptualization: B.L., A.K., and C.M.N. Methodology: B.L., M.R., P.Z., B.W., and S.X.Z. Investigation: B.L., M.R., P.Z., B.W., M.K., S.T., A.S., and K.G. Writing: B.L. and C.M.N. Supervision: E.R.G., R.M., A.K., and C.M.N.

## Data and materials availability

All data needed to evaluate the conclusions in the paper are present in the main text and/or the Supplementary Materials.

## List of Supplementary Materials

Materials and Methods

Figs. S1 to S6

Movie S1, related to Fig. 1: Rotation of 3D model of lung, color-coded for intensity of Tom20.

Movie S2, related to Fig. 1: High-magnification time-lapse fluorescence microscopy of MitoView633 shows mitochondria on the apical side of the cells.

Movie S3, related to Fig. 1: Time-lapse fluorescence microscopy of MitoView633 in embryonic chicken lung, showing an increase in fluorescence corresponding to a nascent epithelial branch forming along the dorsal surface of the epithelium.

Movie S4, related to Fig. 2: High-magnification time-lapse fluorescence microscopy of ATPRed1, showing ATPRed1 outlining the mitochondria located on the apical side of the cells.

Movie S5, related to Fig. 2: Time-lapse fluorescence microscopy of ATPRed1 in cultured lung explant, revealing a progressive rise in ATP levels at future branch sites.

Movie S6, related to Fig. 2: Time-lapse fluorescence microscopy of ATPRed1 in a cultured lung explant treated with blebbistatin, resulting in a drop in ATPRed1 fluorescence.

Movie S7, related to Fig. 3: Brightfield movies of lungs cultured in the presence of vehicle control (DMF) or oligomycin (2 *μ*M).

Movie S8, related to Fig. 4: Finite element simulations with programmed active strains guided by mitochondrial distributions reproduce the morphodynamics of neural tube closure and branch initiation in chicken, optic pit invagination in mouse, and ventral furrow formation in *Drosophila*.

## Supplementary Materials

### Materials and Methods

#### Animal husbandry

All procedures involving animals complied with ethical regulations and were approved by the Princeton University Institutional Animal Care and Use Committee (IACUC). For experiments with embryonic lungs and neural tubes, fertilized chicken (*Gallus gallus* variant *domesticus*) eggs were obtained from the Department of Animal Science, University of Connecticut. Eggs were incubated in an upright configuration at 37°C and 70% humidity for the specified number of hours. Underdeveloped or deformed embryos were discarded. For experiments with embryonic eyes, C57BL6/J mice were bred and maintained under standard laboratory conditions, receiving food and water *ad libitum* in an AAALAC-accredited facility in accordance with the NIH Guide for the Care and Use of Laboratory Animals. Noon of the day on which a vaginal plug was detected was considered as embryonic day 0.5 (*E*0.5). For experiments with gastrulating embryos, *Drosophila melanogaster* strains were reared at 21°C under standard laboratory conditions. For embryo collection, flies were transferred into cages and fed using agarose plates supplemented with apple juice and yeast paste.

#### Wholemount immunofluorescence staining and imaging

Embryonic tissues were isolated and fixed in 4% paraformaldehyde (PFA) with 0.25% glutaraldehyde in phosphate-buffered saline (PBS) with at least a 1:20 ratio of sample volume to fixative volume for 30 min at 4°C on a shaker, followed by an additional 15 min at 20°C on a shaker. Samples were washed in tris-buffered saline (TBS) containing 7.35 mg/mL CaCl_2_ for 15 min on a shaker and then in TBS containing 0.5% Triton X-100 (TBST) three times for 15 min each. Samples were then blocked in blocking buffer comprised of TBST containing 10% donkey serum for 1 hour at 20°C on a shaker. After blocking, samples were incubated in blocking buffer containing primary antibodies against Tom20 (1:100; D8T4N, rabbit mAb; Cell Signaling 42406S), E-cadherin (1:500; clone 36, mouse mAb; BD 610182), or pMLC (1:50; Ser19, mouse mAb; Cell Signaling 3675S) for 48 hours at 4°C. Samples were then washed in TBST four times for 15 min each. Samples were then incubated in blocking buffer containing donkey anti-rabbit (H+L) Alexa Fluor 555 (1:500; ThermoFisher Scientific A31572), donkey anti-mouse (H+L) Alexa Fluor 680 (1:500; ThermoFisher Scientific A10038), or Alexa Fluor 488-conjugated phalloidin (1:500; ThermoFisher Scientific A12379), and then counterstained with DAPI (1:1000; ThermoFisher Scientific D1306). Samples were washed in TBST three times for 15 min each, and then once in PBS for 15 min. Samples were dehydrated in an isopropanol series at 25%, 50%, 75%, and 100% for 15 min each and then titrated into a 1:2 ratio of benzyl alcohol to benzyl benzoate (BA:BB) at 50% and then 100% for 15 min each. Samples cleared in BA:BB were held by a nylon washer attached to two coverslips and imaged using a Hamamatsu MAICO scanning confocal fitted to an inverted Nikon TE microscope with a Ludl stage using a 10× 0.5 *NA* Nikon objective (CFI Super Fluor 10× MRF00101) for whole-organ imaging or a 40× 1.15 *NA* Nikon objective (CFI Apo LWD Lambda 40XC WI MRD77410) for high-magnification imaging.

#### Explant culture and live-imaging analysis

Embryonic lungs were dissected in sterile PBS and transferred to custom-made imaging chambers consisting of a #1.5 coverslip coated with a thin layer of polyacrylamide (**Fig. S6**). Samples were cultured under a thin layer of DMEM/F12 medium (without HEPES or Phenol Red) supplemented with 5% fetal bovine serum (FBS, heat-inactivated; Atlanta Biologicals) and antibiotics (50 units/mL of penicillin and streptomycin). A 1-mL reservoir of culture media was created by surrounding the explants in a ring of hydrated, sterilized, fibrous material. Timelapse imaging was performed within an OKO stage-top incubator at 37°C, 5% CO_2_, and 75% relative humidity. Confocal fluorescence images were acquired using a Hamamatsu MAICO scanning confocal fitted to an inverted Nikon TiE microscope with a Ludl stage and a 20× 0.95 *NA* WI Nikon objective (CFI Apo LWD Lambda S 20XC WI MRD77200). Phase-contrast images were acquired using a 4× 0.2 *NA* objective (CFI Plan Apochromat Lambda D 4X MRD70040); a green filter was placed on the brightfield arm at the condenser to reduce damage from light.

Stock concentrations of MitoView633 (200 mM; Biotium #70055) and ATPRed1 (4.5 mM; Sigma-Aldrich SCT045) were prepared in DMF. To prevent the precipitation of ATPRed1, we generated a final stock of 1.4-mM ATPRed1 by mixing 4 μL of PEG300 (ThermoFisher Scientific 192220010) with 2 μL of prepared dye, followed by adding 0.5 μL of Tween20. Stock mixtures were sonicated for 2 min after preparation and again for 2 min immediately before use. We used pulled-glass needles (<5 μm in diameter) to create holes in the distal tip of the embryonic lung epithelium. We then loaded 40-μM MitoView633 or 1.1-mM ATPRed1 into pulled-glass needles and injected ∼0.1 *μ*L into the carina of the lung for live-imaging analysis (**Fig. S6**). Fluorescence images were acquired 30 min after injection.

For experiments with gastrulating embryos, *Drosophila melanogaster* embryos were collected at the cellularization stage, dechorionated, mounted on a coverslip using heptane glue, and covered with Halocarbon Oil 200 before imaging.

For live-imaging analysis of the neural tube, fertile eggs were incubated for 32 hours such that embryos developed to stage HH8-9. Embryos were collected on filter paper using the EC protocol (*36*) and treated with ATPRed1 and MitoView633 diluted in media (1:1 for ATPRed1; 1:8 for MitoView633), added to the embryonic ectoderm using a 20-*μ*L pipette, locally mixing the dye along the neural tube. Embryos on filter paper were then cultured dorsal side down in a glass-bottom dish in egg albumin at 37°C for live-imaging experiments.

For live-imaging analysis of embryonic mouse eye placodes, embryos were collected at *E*10.5 and placed into glass-bottom plates containing media [DMEM with sodium pyruvate (100 mM), HEPES (1 M), MEM non-essential amino acids, penicillin-streptomycin, and 10% FBS]. ATPRed1 and MitoView633 were first diluted in media (1:1 for ATPRed1; 1:8 for Mitoview633). The embryo was placed on its side, then 10 µl of the diluted dye mixture was pipetted onto the eye placode. The embryo was then flipped so that the eye placode labeled with ATPRed1 and MitoView633 would be in contact with the glass-bottom plate for live-imaging analysis.

To inhibit actomyosin contraction or mitochondrial ATP generation, lung explants were cultured in media containing 1:1000 para-amino-blebbistatin (Cayman #22699) or oligomycin (Sigma-Aldrich #75351) dissolved in DMF at the concentrations indicated. As a vehicle control, lung explants were cultured in media containing 1:1000 DMF.

#### Virus production and injection

The PercevalHR RCAS expression construct was assembled by subcloning the PercevalHR insert from GW1-PercevalHR (Addgene #49082) into an RACS(BP) avian retroviral backbone. The recombinant RCAS plasmid was transfected into chick DF1 cells in T-75 flasks using standard transfection protocol. The transfected cells were further maintained in T-175 flasks until confluency. Cell culture medium was harvested from confluent populations once per day for four days, and concentrated at 45,000 rpm for 1.5 hours. The pellet was dissolved in minimal volume of viral resuspension buffer (20-mM Tris pH 8.0, 250-mM NaCl, 10-mM MgCl_2_, 5% sorbitol) and stored at -80°C until use. For introduction of PercevalHR RCAS into chicken embryos, fertile eggs were incubated for 32 hours such that embryos developed to HH8-9, followed by windowing of the egg and injection of concentrated virus into the coelomic cavity. Windowed eggs were resealed and incubated for an additional four days, after which time lungs were dissected and analyzed using confocal microscopy. PercevalHR images were acquired on a Nikon AXR scanning confocal microscope using a 20× 0.75 *NA* objective (CFI SuperFluor 20X MRF00200), imaging separately first with a 488-nm laser for *λ*_high_ and then with a 405-nm laser for *λ*_low_, and collected with a 530/40 filter. The final ratiometric signal is a 32-bit division of *λ*_high_/*λ*_low_.

#### Image analysis

For fixed samples imaged at 10×, epithelial and mesenchymal regions were segmented in ImageJ by applying a threshold to the E-cadherin (Ecad) channel. Custom Julia code (available at github.com/bezlemma) was then used to remove the esophagus from the image. In cases where the esophageal signal was very close to that of the lung, Ecad signal was manually deleted in ImageJ. The original confocal volume data and associated segmentations are provided alongside the code.

Within these segmented regions, the mean Tom20 signal intensity was calculated separately for the epithelium and mesenchyme. To define proximal and medial sections, y-coordinates were used to identify the carina and the first branch; data between the carina and first branch were defined as the proximal region. Data halfway between the last visible branch and the distal tip were defined as the distal region. Data between these two regions were defined as the medial region. Julia code for both segmentation and mean-intensity calculations (with y-coordinates) are included in the repository.

For both fixed and live samples imaged at 20×, manual measurements were performed to calculate the ratio of fluorescence intensities—Tom20 (mitochondrial density), MitoView633 (mitochondrial potential), or ATPRed1 (mitochondrial ATP)—between the dorsal and ventral epithelium. These measurements were made in four specified regions by using the ImageJ lasso selection tool.

In continuous temporal confocal datasets, the images were first stabilized in ImageJ. A square selection was placed over the epithelial area of interest at each time point, and the mean intensity within this selection was recorded. For continuous spatial data, a freehand line tool with a width corresponding to the approximate thickness of the epithelium (∼30 μm) was used to measure mean intensity across that width. All datasets were then manually aligned so that branching epithelial regions overlapped. The raw data from these manual steps are included within the plotting scripts provided in the code repository.

#### Spatial transcriptomic mapping

We performed spatial transcriptomic mapping using a modified microfluidic-enabled, deterministic barcoding-based (DBiT-seq) workflow (*37*). Briefly, lungs from 124-hour-old embryos were embedded and sectioned into 7–9-µm-thick slices using a cryostat, then mounted at the center of poly-L-lysine-coated glass slides. The sections underwent spatial barcoding through two rounds of barcode application. In the first round, barcodes A1 to A50 were introduced into the tissue sections via 15-µm-wide microfluidic channels, followed by in situ reverse transcription. In the second round, barcodes B1 to B50 were applied orthogonally to the first set using another microfluidic device, accompanied by in situ ligation. The barcoded tissue sections were subsequently digested, and the resulting cDNA was purified and sequenced according to established protocols (*38*).

#### Oxygen consumption rate measurements

Lung explants were cultured at 37°C and 90% humidity in Agilent Seahorse XF DMEM, pH 7.4 (Agilent #103575-100), supplemented with 5% FBS, antibiotics (50 units/mL of penicillin and streptomycin), glucose (10 mM; Agilent #103577-100), pyruvate (1 mM; Agilent #103578-100), and glutamine (2 mM; Agilent #103579-100). Prior to culture, Islet Capture Microplates (Agilent #101122-100) were coated with a central circle of 2-μL poly-D-lysine. Freshly dissected lung explants were placed in the middle of the circle, submerged in 500-μL media, and then covered with nylon islet capture screens. Images of each explant were acquired using an Olympus SZX7 stereoscope fitted with an OMAX A35140U camera to permit normalization of the data by projected area of the tissue. Data were discarded from any wells in which lungs had been displaced out of the central circle during the process. Oxygen consumption rates were measured using an Agilent Seahorse XFe24 Analyzer following the manufacturer’s protocols; pharmacological agents were injected at the concentrations indicated, with 56 μL in Port 1 and 62 μL in Port 2.

#### Reaction-diffusion model of ATP hydrolysis

We model the concentrations of [ATP] and [ADP] in a 1D domain [0, L] that represents the apicobasal axis of the cell. Each chemical species satisfies a diffusion equation with spatially dependent reaction terms

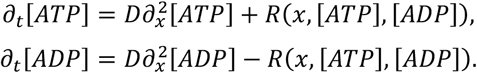

Using finite difference, we numerically calculate the density of chemical species A as

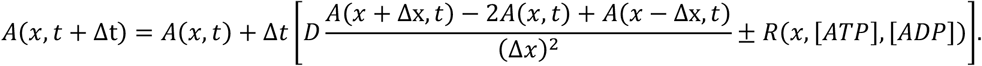

At the boundaries we impose a no-flux boundary condition:

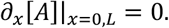

This condition is implemented numerically by modifying the diffusion term at the boundaries to

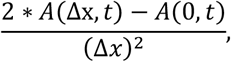

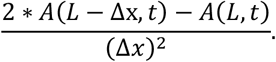

The reaction term R includes a baseline conversion of ATP to ADP everywhere, representing the cell’s baseline ATP usage, *R*_0_. The reaction term also includes an ADP-to-ATP conversion (phosphorylation), *R*^*ADP*→*ATP*^, which applies proportionally to a mitochondrial density function *M*(*x*). Finally, the reaction term includes an ATP-to-ADP conversion term (hydrolysis), *R*^*ATP*→*ADP*^, which applies proportionally to actomyosin density Θ(*x*):

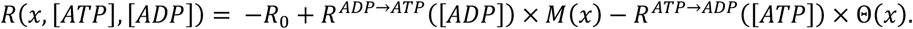

The phosphorylation term is such that no timestep phosphorylates more ADP than is available.

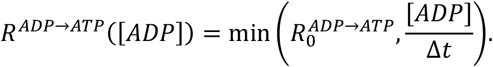

The hydrolysis term has a Michaelis-Menten-like dependence, with [ADP] as an additional inhibitor with *K*_*i*_ = *K*_*m*_, such that

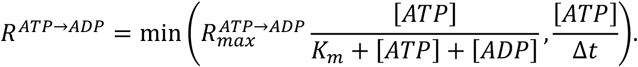

Where 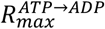 is the maximum hydrolysis rate, *K*_m_ is the Michaelis constant, and a minimum function is used to ensure that enough ATP exists for the step to take place. 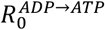 is calculated from experimental measurements of mitochondrial respiration. Simulations are shown with 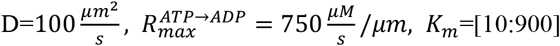, and 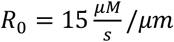.

#### Estimation of ATP phosphorylation rate from experimental measurements of oxygen consumption rate

To estimate the rate of ATP production from mitochondria, *∂*_*t*_[*ATP*]_*mito*_, from the total rate of oxygen consumption, *∂*_*t*_[*O*_2_], we measured the rate of oxygen consumption when the ATP-producing complex V of mitochondria was inhibited by treatment with oligomycin, *∂*_*t*_[*O*_2_]_*other*_. This term, *∂*_*t*_[*O*_2_]_*other*_, encompasses both the mitochondrial proton leak as well as oxygen consumption due to non-mitochondria-related events. Thus, the oxygen consumption rate due to mitochondria is

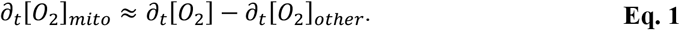

Since there are approximately 5.5 molecules of ATP produced per molecule of O_2_ consumed (*39*), we calculate mitochondrial ATP production as

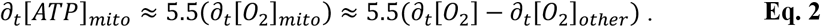

The energy produced by phosphorylating ADP into ATP, *E*_*ADP*→*ATP*_, has both a constant term,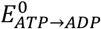, and an entropic term dependent on the logarithm of the ADP/ATP ratio. Specifically,

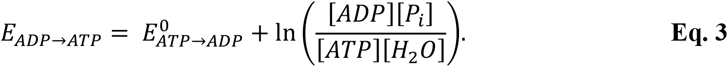

To estimate the chemical energy produced by mitochondria from oxygen consumption rates, we take the ADP/ATP ratio such that *E*_*ADP*→*ATP*_ ≈ 10^−19^*J* uniformly (*27*). We convert this energy into power, or energy per unit time:

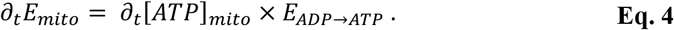

The resulting estimated chemical power of ATP phosphorylation produced by mitochondria, *∂*_*t*_[*ATP*]_*mito*_, is ∼ 60 *μJ*/*min* or ∼ 1*μW*. Over several hours this accounts for a chemical energy that is 100-million-fold the mechanical deformation energy derived below.

#### Calculation of viscoelastic energy requirements for the initiation of a branch by apical constriction

Viscoelastic energy change from change in volume, Δ*E*_*stretch*_, has an elastic energy density term plus a viscous dissipation term:

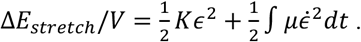

Where *K* is the bulk modulus, *ϵ* is the volumetric strain, and *μ* is the viscosity. Over time, Δ*t*, this becomes

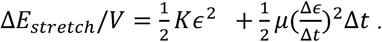

Consider the deformation of a half-sphere with radius *R*_1_ from a flat sheet as a proxy for the deformations studied here:

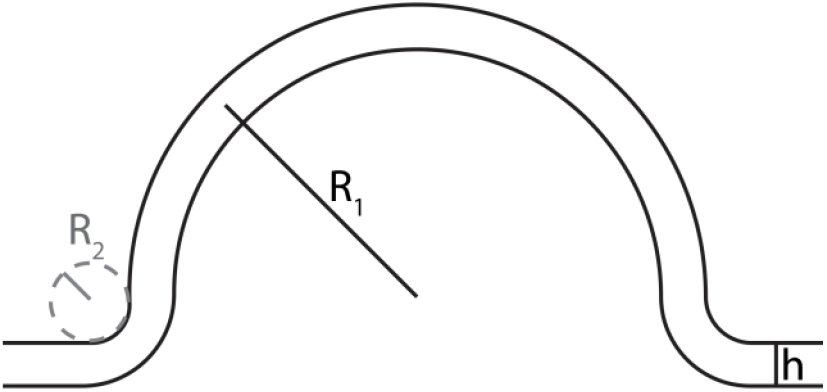

Note that the volumetric strain, 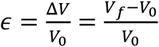, is *ϵ* = 1, as a circular patch with height *h* and radius *R*_1_ gives 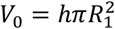, while deformation into a half sphere gives 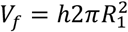. Thus,

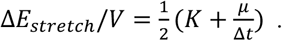

Taking physical constants as *K* ≈ 10 *kPa*, Δ*t* ≈ 4 *hrs, μ* ≈ 10 *Pa* ⋅ *s, h* = 30 *μm*, and *R*_1_ = 100 *μm*, we find that Δ*E*_*stretch*_ ≈ 5 *nJ*.

To calculate the change in energy due to bending, we assume a form of local bending energy density with principal curvatures *k*_1_ and *k*_2_ to be

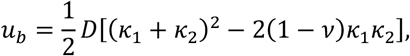

where 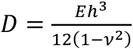 is the flexural rigidity. Taking this for our sphere, where 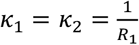, we integrate the local bending energy density of a thin sheet with curvature *R*_1_,

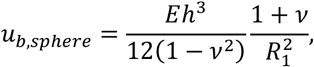

over the area of the formed half sphere

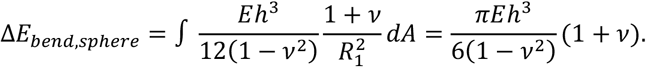

Then considering the neck region, where 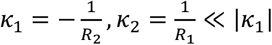, the local bending energy density is approximately

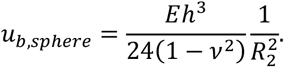

By integrating over the neck region with area ≈ 2*πR*_1_*R*_2_ we obtain the bending energy of the neck as

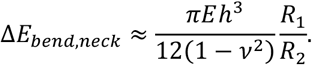

The total bending energy is thus

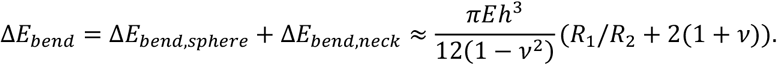

with radius *R*_1_ ≈ 100 *μm*, and a curvature of *R*_2_ ≈ 5 *μm*, Young’s modulus *E* ≈ 10 *kPa*, thickness of the epithelium *h* ≈ 30 *μ*m, and Poisson ratio *v* ≈ 0.3. These parameters result in the bending energy Δ*E*_*bend*_ ≈ 2 *nJ*.

Thus, to an order of magnitude, the total energy due to this idealized deformation is

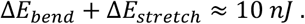

#### Computational model

The 3D simulations of epithelial morphogenesis in response to apical constriction use a compressible Neo-Hookean hyperelastic constitutive law, solved via the finite element method (FEM). We assumed that actomyosin-driven apical constriction is much slower than mechanical relaxation and, hence, the system is always in quasi-mechanical equilibrium, which was achieved by minimizing the total elastic energy as described below.

Supposing that the initial reference volume is Ω, we introduced a fixed Cartesian coordinate system with an orthonormal basis {***e***_**1**_, ***e***_**2**_, ***e***_**3**_} and spatial coordinates given as (*X*_1_, *X*_2_, *X*_3_), denoted as ***X*** = *X*_*i*_***e***_***i***_ where summation over repeated indices is implied. At some later time *t*, the system was deformed to a volume Ω_*t*_ and the Cartesian coordinates were mapped to a different vector field, denoted as ***x*** = φ_*t*_(***X***). Using this notation, we follow finite deformation theory (*40*) and define the deformation gradient tensor as 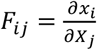.

To implement apical constriction, we decompose the total deformation gradient ***F***(***X***) as ***F*** = ***F***_***e***_***F***_***c***_, where it is assumed that isotropic contractions induce an intermediate stress-free state with deformation gradient ***F***_***c***_ = (1 − *tβ*)***I***, where *β*(***X***) is the contractility field and *t* is time, and that there is an additional elastic deformation of this intermediate stress-free state to the final deformed state due to the elastic deformation gradient ***F***_***e***_. The total deformation gradient can be expressed in terms of the displacement field ***u***(***X***) = ***x***(***X***) − ***X*** as ***F*** = ***I*** + ∇***u***. Thus, the elastic deformation gradient can be expressed as ***F***_***e***_ = ***FF***_***c***_^−1^ = (1 − *tβ*)^−1^(***I*** + ∇***u***).

To account for large material deformation, we assumed the tissues to be Neo-Hookean solids with the elastic energy storage density *ψ*(***F***_***e***_, ***X***) defined as

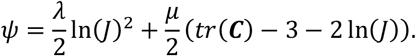

In which λ(***X***) and μ(***X***) are the Lamé constants, ***C*** is the Cauchy-Green deformation tensor, and *J* is the Jacobian of the elastic deformation gradient ***F***_***e***_. Explicitly,

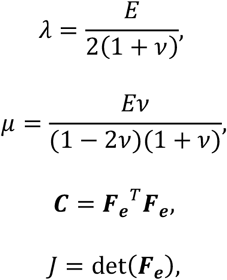

where ν is the Poisson ratio and *E* is the elastic modulus.

The FEM, a standard method in computational mechanics to numerically solve for the deformation field under given boundary conditions (*41*), was used to calculate the displacement field ***u***(***X***) that minimizes the total potential energy Π(***u, λ***_*tr*_, ***λ***_*rot*_) for a prescribed contraction profile where the Lagrange multipliers ***λ***_*tr*_and ***λ***_*rot*_ constrain rigid body translation and rotation, respectively. The total potential energy can be written as (*42*):

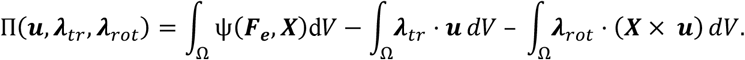

In all simulations we use traction-free boundaries. For the neural tube geometry (**Fig. S5b,d**), a boundary condition enforces that the z-displacement at top-z of the geometry is 0, preventing the planar parts of the geometry from folding upwards. The unknown displacement field ***u*** is then obtained via the variation of the potential energy function Π(***u, λ***_*tr*_, ***λ***_*rot*_) with respect to ***u, λ***_*tr*_, and ***λ***_*rot*_ and solving,

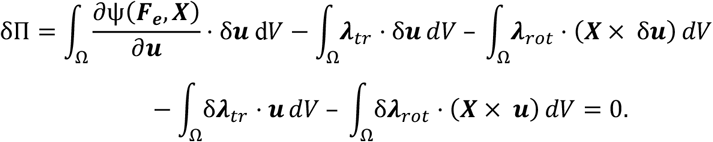

For each timestep, we solve displacements ***u***(***X***) by minimizing the total elastic energy Π using the Newton-Raphson method, then the next timestep occurs with an increase in contraction according to *β*(***X***) and the process is repeated. If the Newton-Raphson method fails to converge, we reduce the timestep and repeat.

To numerically solve the above energy minimization problem, the domain Ω was discretized using first-order tetrahedral elements that were generated with the help of an open-source software Gmsh (*43*). Then, the potential energy minimization problem was implemented in the open-source computing platform FEniCS (*44*). Simulations contained ∼20,000 tetrahedral elements, chosen by serial refinement until simulation results remained consistent. Geometrical parameters were chosen to match confocal imaging data. The non-dimensional elastic modulus was defined as *E* = 1 and the Poisson ratio was set as ν=0.3. All simulation code and geometries can be found at github.com/bezlemma. Simulations were visualized in ParaView (*45*).

## Supplementary Figures

**Fig. S1.**
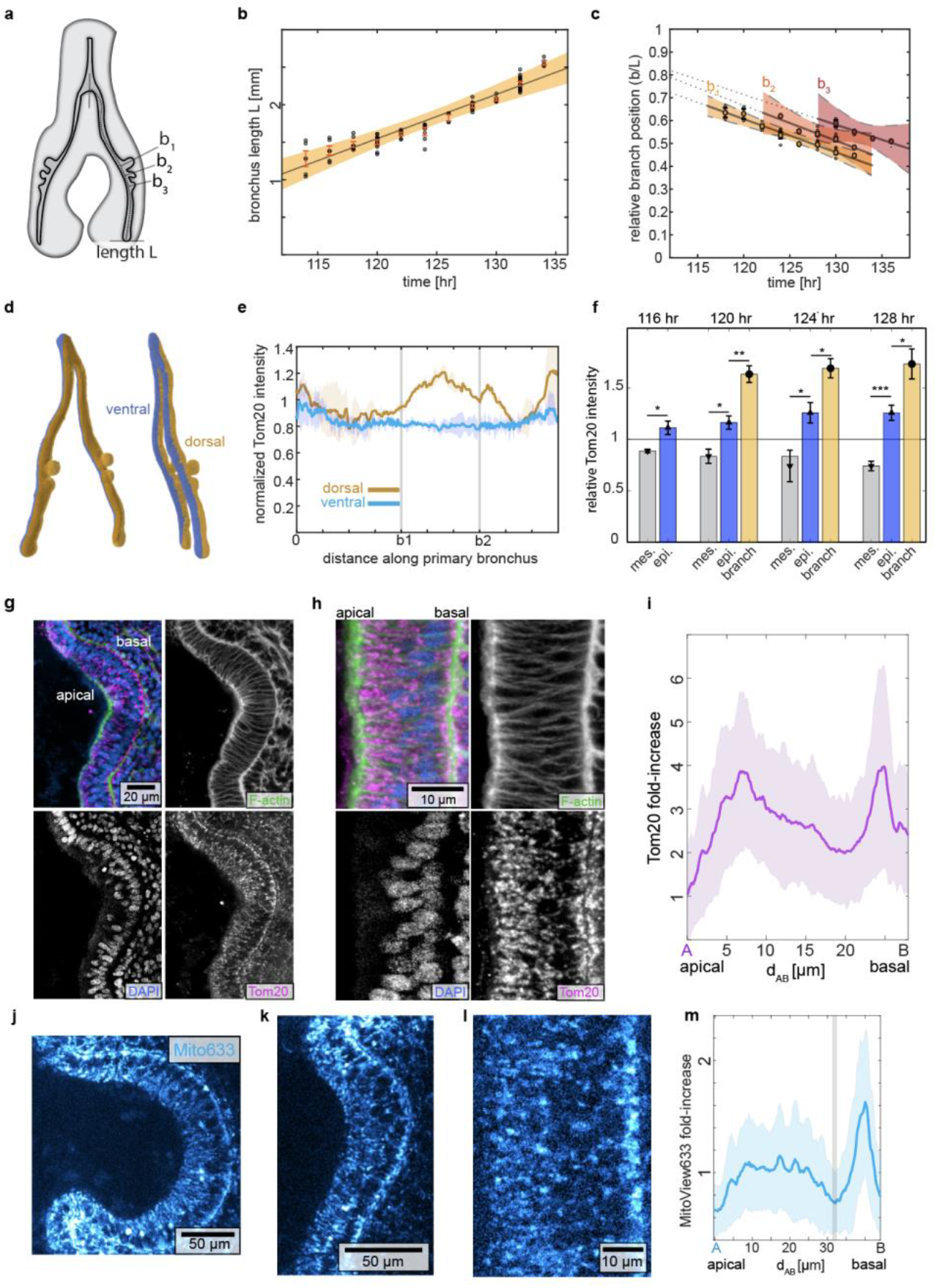
Characterization of mitochondrial membrane density and potential in the early embryonic chicken lung – related to Fig. 1. (**a**) Schematic of HH26-stage lung, showing first branch (b1), second branch (b2), third branch (b3), and length of the primary bronchus (L). (**b**) Graph showing L as a function of developmental time. Each smaller dot represents one lung. Shaded region represents the 95% confidence interval of the linear fit. (**c**) Graph showing the relative positions of b1, b2, and b3 as a function of developmental time, normalized to L. Shown are data from 6 lungs for each timepoint. Shaded region represents the 95% confidence interval of the linear fits. (**d**) Volume rendering of epithelium within HH26-stage lung, color-coded to indicate dorsal (gold) and ventral (blue) regions of the epithelium. (**e**) Graph of normalized Tom20 staining intensity in the dorsal (gold) and ventral (blue) epithelium, as a function of distance along the primary bronchus, normalized such that 1 is the mean Tom20 intensity of the lung. (**f**) Graph of Tom20 staining intensity in the mesenchyme, total epithelium, and branching epithelium as a function of developmental stage. Error bars indicate s.e.m. for 3 lungs at the 0-branch stage (116 hr) and 5 lungs at each subsequent stage. (*) indicates p<0.051, (**) p<0.01, (***) p<0.001, (**g**) Fluorescence images of F-actin (green), Tom20 (magenta), and nuclei (blue) in a single branch of a developing lung. Scale bar, 20 *μ*m. (**h**) Fluorescence images of staining for F-actin (green), Tom20 (magenta), and nuclei (blue) at an uncurved future branch site in the dorsal epithelium. Scale bar, 10 μm. (**i**) Graph showing quantification of normalized Tom20 intensity as a function of position along the apicobasal axis, averaged along the proximal-distal axis of a dorsal branching region. Shaded error bar indicates standard deviation along the proximal-distal axis. Fluorescence images of MitoView633 dye visualizing mitochondrial membrane potential in (**j**) a branch, (**k**) a nascent branch, and (**l**) an uncurved future branch site in the dorsal epithelium. Scale bars, 50, 50, 10 *μ*m. (**m**) Graph showing quantification of normalized MitoView633 intensity along the apicobasal axis of a branching region. Shaded error bar indicates standard deviation of integrated intensity.

**Fig. S2.**
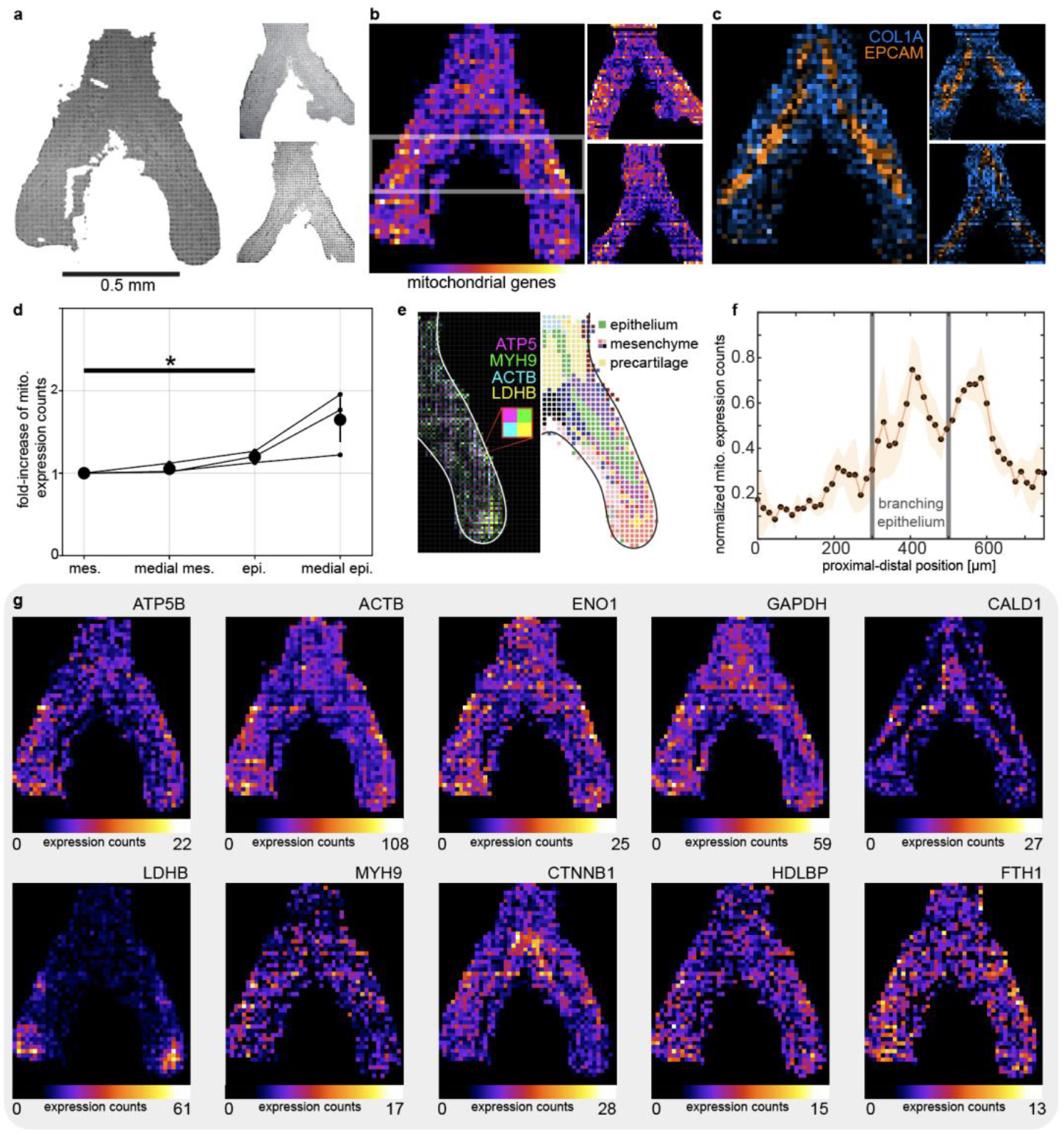
Unbiased spatial transcriptomics reveals increased expression of mitochondrial genes in the branching lung epithelium – related to Fig. 1. (**a**) Segmented images of the three lung slices used for DBiTseq. (**b**) Merged expression counts of mitochondrial genes; white bars bracket the area considered as medial for later analysis. (**c**) Maps of COL1A and EPCAM expression used to segment the mesenchyme and epithelium, respectively. (**d**) Graph indicating fold increase of mitochondrial gene counts, as compared to the mesenchyme. Data show significant difference (p=0.03) in mitochondrial gene expression between the mesenchyme and epithelium. (**e**) Composite map of genes encoding for ATP synthase (ATP5), non-muscle myosin heavy chain (MYH9), beta-actin (ACTB), and lactate dehydrogenase b (LDHB). (**f**) Graph showing intensity of mitochondrial gene expression along the proximal-distal axis of the dorsal epithelium, defined as the outer third of the tissue, from carina to distal tip, revealing higher mitochondrial gene-expression counts in the branching and future branching regions of the lung. Shaded region is s.e.m. across three independent replicates. (**g**) Spatial maps of expression counts of ATP synthase F1*β* subunit (ATP5B), ACTB, enolase-1 (ENO1), glyceraldehyde 3-phosphate dehydrogenase (GAPDH), caldesmon (CALD1), LDHB, MYH9, beta-catenin (CTNNB1), high-density lipoprotein binding protein (HDLBP), and ferritin heavy chain (FTH1).

**Fig. S3.**
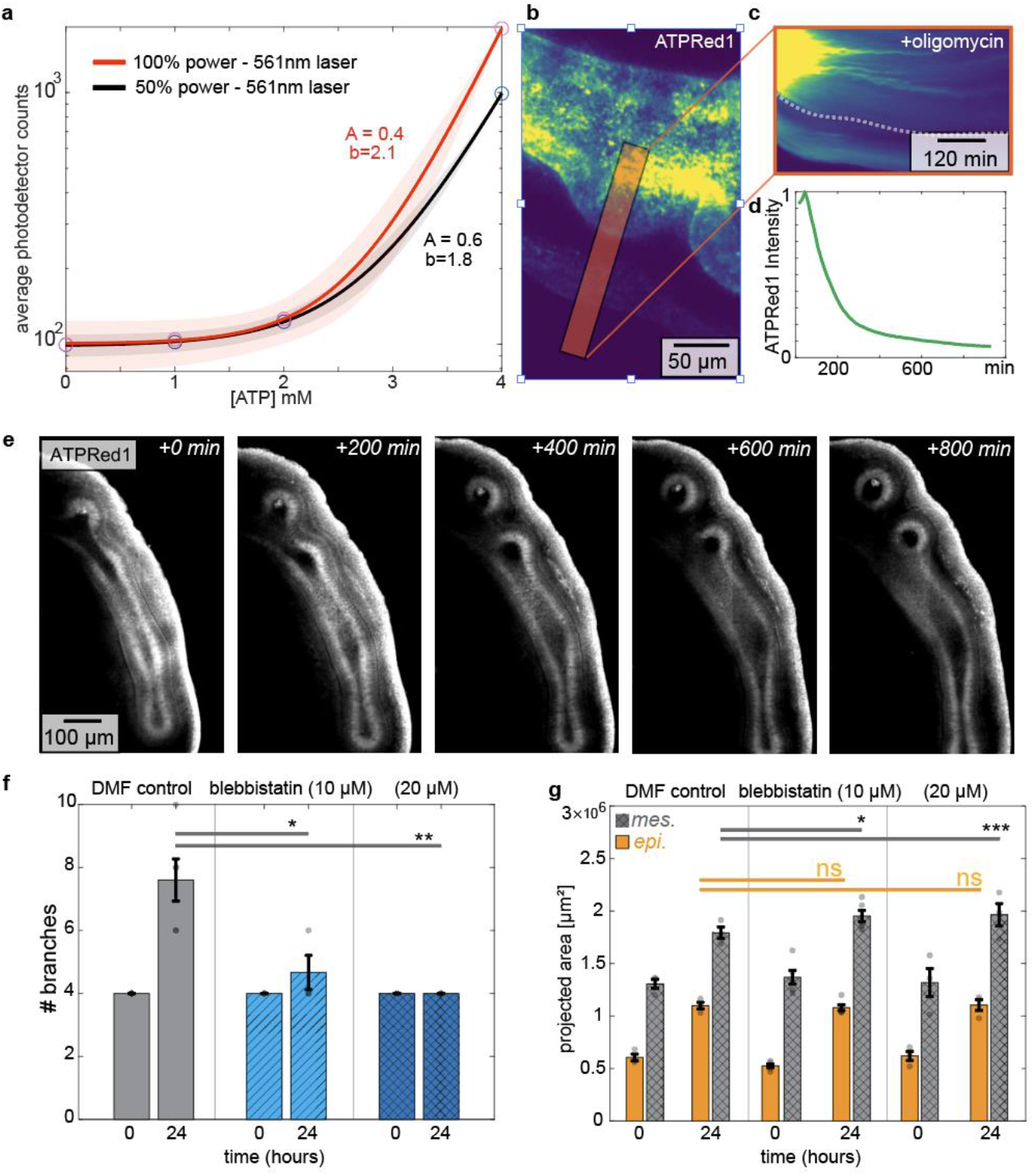
Characterization of mitochondrial ATP and actomyosin contractility in the developing lung – related to Fig. 2. (**a**) Graph showing calibration of ATPRed1 fluorescence intensity as a function of ATP concentration [ATP] for two different laser intensities. Lines show the best exponential fits for the data. Shaded error bar indicates the 95% confidence interval of the fit. (**b**) Fluorescence image of ATPRed1 labeling of lung explant. Scale bar, 50 μm. (**c**) Kymograph of ATPRed1 fluorescence intensity as a function of time after treatment with oligomycin (2 μM), for region indicated in (b). Time bar, 120 min. (**d**) Graph showing change in relative ATPRed1 fluorescence intensity as a function of time after treatment with oligomycin. (**e**) Timelapse fluorescence images of z-projected average ATPRed1 intensity in embryonic lung explant. A single z-slice is shown in Movie S5. Scale bar, 100 μm. (**f**) Graph showing number of branches as a function of time in lung explants cultured in the presence of DMF control or blebbistatin (10 μM or 20 μM). Error bars indicate s.e.m. for 3 explants across 3 independent experiments. *, ** indicate p=0.04, p=0.004. (**g**) Graph showing projected area of the epithelium and mesenchyme as a function of time in lung explants cultured in the presence of DMF control or blebbistatin (10 μM or 20 μM). Error bars indicate s.e.m. for 6 explants across 3 independent experiments. *, ** indicate p=0.04, p=0.001.

**Fig. S4.**
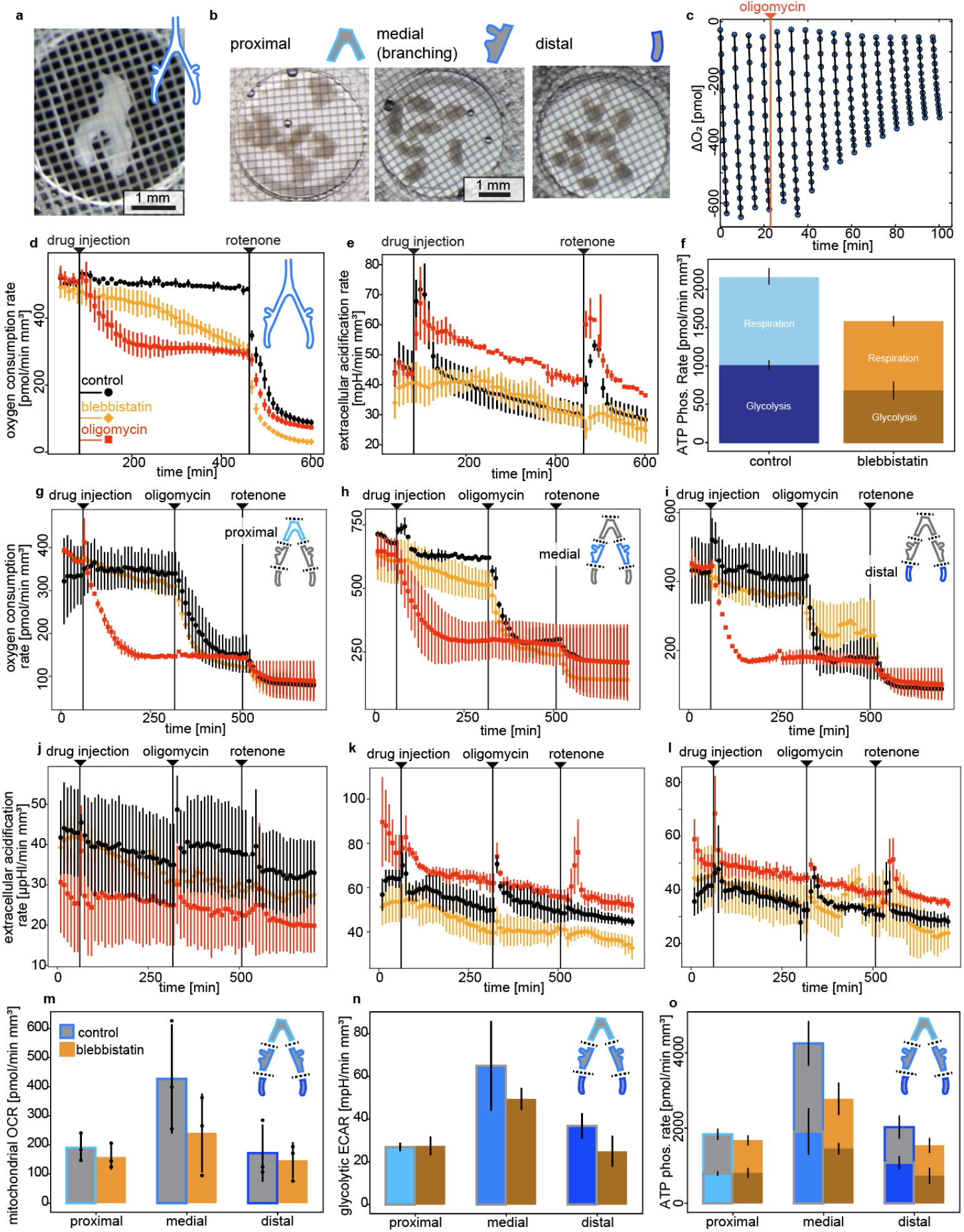
Characterization of oxygen consumption rate and extracellular acidification rate in embryonic lung explants – related to Fig. 2. (**a**) Brightfield image of an intact lung placed in a Seahorse XFe24 Islet Well with a containing grid on top of it. Scale bar, 1 mm. (**b**) Brightfield images of proximal, medial, and distal portions of lungs pooled in XFe24 Islet Wells. Scale bar, 1 mm. (**c**) Graph showing an example raw measurement of oxygen levels. Between each line, vibrations reoxygenate the media in the well before the next series of measurements is taken. Dots represent oxygen readings, while the slopes of the line fits are the oxygen consumption rates. (**d**) Graph showing oxygen consumption rates for intact lungs treated with oligomycin, blebbistatin, or DMF control, followed by a rotenone injection. Error bars are the standard error over 3 experimental replicates; each replicate contains 16 lungs each measured in a separate well. (**e**) Graph showing extracellular acidification rates for the same data sets as in (d). (**f**) Graph showing calculated ATP phosphorylation rate due to mitochondrial respiration and glycolysis for intact lungs and lungs exposed to blebbistatin. Error bars are standard error over the experimental replicates. (**g-i**) Graphs showing oxygen consumption rates for pools of proximal, medial, and distal portions of 24 lungs. Error bars indicate standard error from 3 experimental replicates. (**j-l**) Graphs showing extracellular acidification rates for the same data sets as in (g-i). (**m**) Graph showing mitochondrial oxygen consumption rates calculated from the difference between the control/blebbistatin measurements and the oligomycin measurements. (**n**) Graph showing glycolytic extracellular acidification rates calculated from the value of control/blebbistatin rates after rotenone injection. (**o**) Graph showing the extrapolated ATP phosphorylation rate due to mitochondrial respiration and glycolysis for intact lungs and lungs exposed to blebbistatin. Error bars represent s.e.m.

**Fig. S5.**
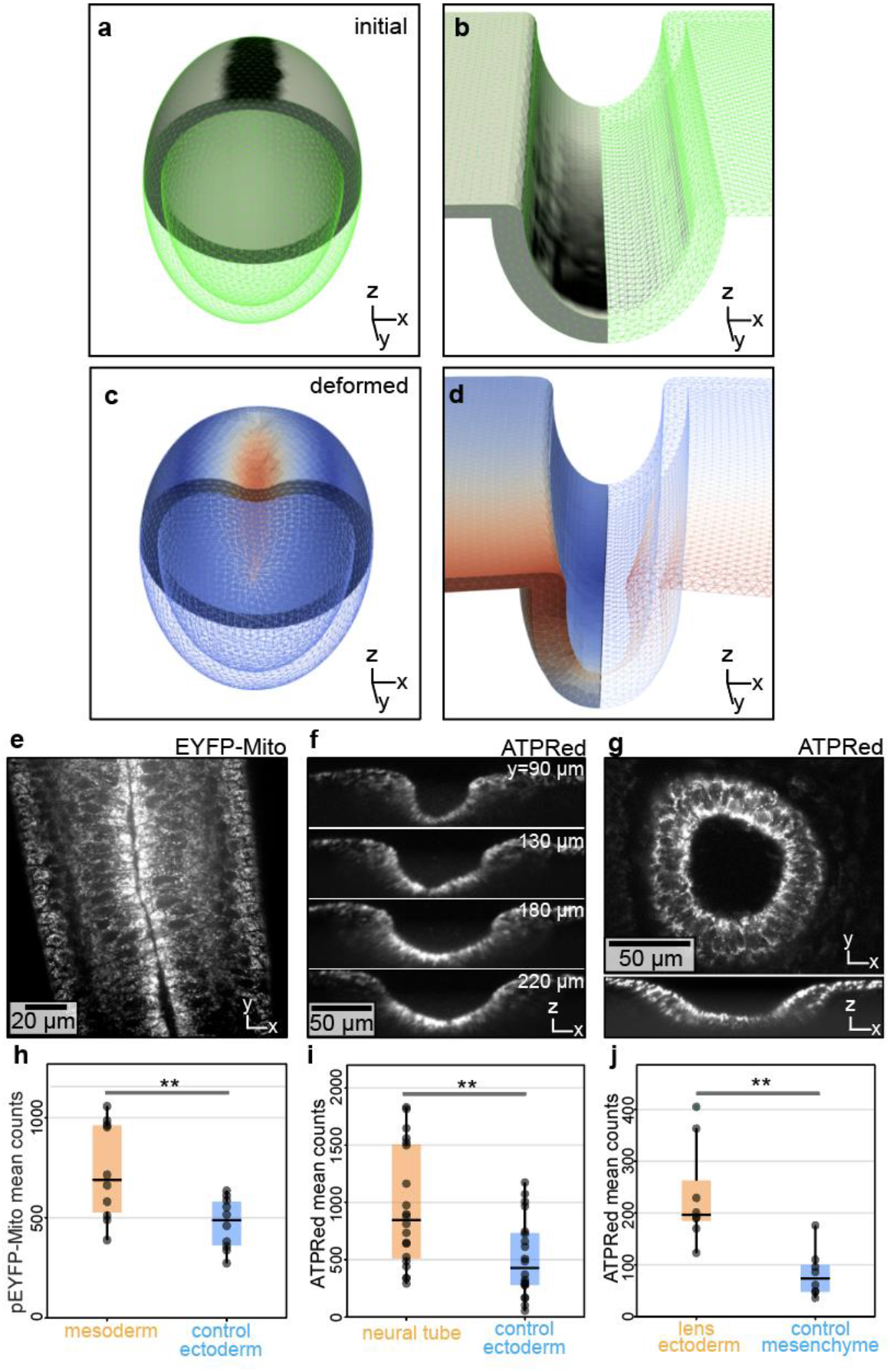
Active strain simulations of apical constriction across species and tissues correspond to apicobasal patterns of enriched mitochondria – related to Fig. 4. (**a-b)** Cutaway view of initial geometries for simulations based on mitochondrial patterning of deformations during formation of the ventral furrow and neural tube; numerical mesh drawn in green, with applied active stress field shown in grayscale. (**c-d**) Cutaway view of numerical meshes deformed by active stresses from (a-b), colored by the displacement field. Live imaging of (**e**) pEYFP-mito intensity in *Drosophila* ventral furrow and ATPRed1 in (**f**) chick neural tube or (**g**) mouse lens placode. (**h-j**) Graphs showing mean intensity of pEYFP-mito intensity (p=0.0098) or ATPRed1 (p=0.0035, p=0.0027) as a function of position in each tissue.

**Fig. S6.**
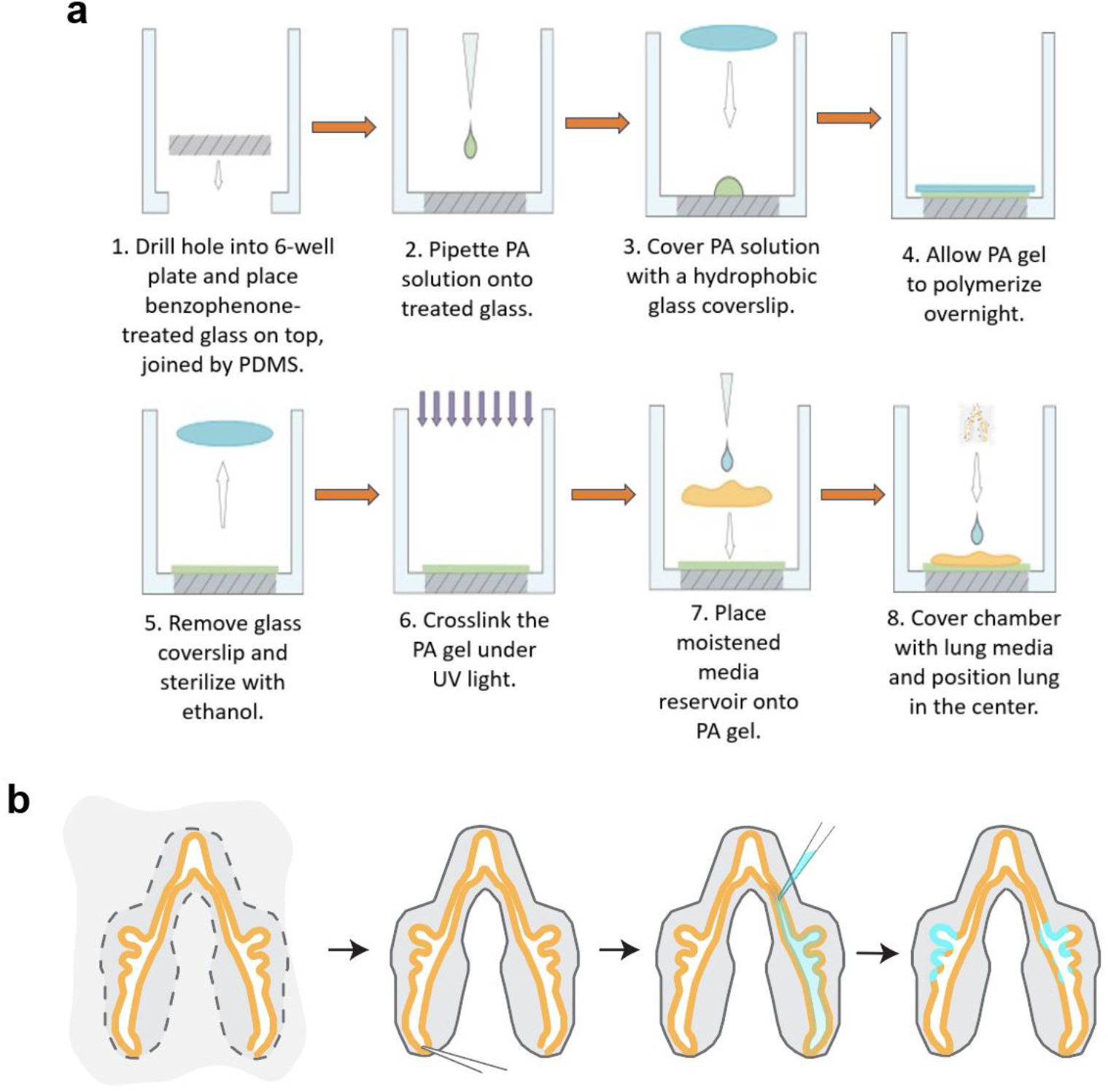
Schematics of live-imaging setup. (**a**) Schematic illustrating construction of chambers for live imaging lungs at high NA. (**b**) Steps for injecting dye into the lumen of the chicken lung. First, the embryonic lung is resected from the embryo. Second, a pulled-glass pipette is used to make small openings at the distal tips to facilitate fluid flow. Third, dye is introduced near the trachea. Finally, the dye passes from the lumen into the epithelial cells.

